# The number of effectors limits the relative sleep gain: Insights from seven finger tapping experiments

**DOI:** 10.1101/2024.05.15.593197

**Authors:** Daniel Erlacher, Daniel Schmid

## Abstract

The role of sleep in motor memory consolidation is still a matter of ongoing debate. A classic task to investigate mechanisms of motor memory consolidation is the finger tapping task, which reliably shows small effects in performance enhancement after sleep but not after a corresponding wake interval. However, variants of the task with a varying number of effectors (e.g., one hand) failed to demonstrate this effect on motor memory consolidation. Thus, in a series of seven experiments we investigate five variants of the classic finger tapping task in which the number of effectors (1 or 2 hands combined with 1, 2 or 4 fingers) used to perform the task are systematically varied. For the groups, where sleep immediately followed learning, a beneficial effect of sleep in comparison with a corresponding wake interval was found, except for the task variant where the finger tapping task was performed with 1 hand and 1 finger. However, no clear-cut pattern could be identified for the numbers of effectors used to perform the task and the magnitude of offline motor memory consolidation. Furthermore, for groups with an intervening wake interval between learning and sleep no differences between the post-sleep and post-wake gain were observed.

**Highlights:** In a variation of the classical finger tapping task with one hand and one finger, no sleep-dependent enhancement was found.

In all other variations, small to large effects of sleep-dependent offline-gains were found.

An interposed wake interval between learning and sleep substantially diminishes the post-sleep enhancement.

Motor skill complexity, with respect to the coordination of more than one effector, does partially play a role in predicting sleep-dependent motor memory enhancement.

## 1 Introduction

The role of sleep in motor memory consolidation is still a matter of ongoing debate (King, Hoedlmoser, Hirschauer, Dolfen, & Albouy, 2017). With an influential theory, Walker (2005) proposed that sleep induces an offline improvement in motor performance when compared with wakefulness (‘Consolidation-based enhancement during sleep’). The idea behind this theory is that sleep-dependent fine-tuning of task-relevant neural circuits are thought to reinforce recently acquired and initially unstable memory traces in an active process (Walker, Stickgold, Alsop, Gaab, & Schlaug, 2005). The most often used task to demonstrate the sleep-dependent consolidation is the classic finger tapping task (FTT), in which an explicitly known sequence (e.g., 4-1-3-2-4) must be repeatedly pressed on a standard keyboard as fast and as accurately as possible. In the first studies using this task in the context of sleep, a night of sleep resulted in about a 20% increase in motor speed without loss of accuracy; whereas the same time spent awake did not lead to any significant improvement (Walker, Brakefield, Morgan, Hobson, & Stickgold, 2002; Walker, Brakefield, Seidman, et al., 2003).

The beneficial effect of sleep in comparison with a wake interval has been repeatedly replicated using an FTT in healthy adults (Schmid, Erlacher, Klostermann, Kredel, & Hossner, 2020). However, few studies could not replicate these findings with a whole night of sleep (Cai & Rickard, 2009), a daytime sleep period (Landry, Anderson, & Conduit, 2016) benefits were only observed with respect to subsequent on-task performance (Maier et al., 2017). Yet as the post-sleep gain in the FTT is exceedingly consistent, the question arises what variations of the original task would limit offline gains and therefore possibly explain the mechanism of motor memory consolidation. One observation from our research has been that no beneficial effect of sleep can be found for a gross motor variant of the classic task, wherein the FTT sequence is performed with(Erlacher, Schmidt, & Blischke, 2014).

Thus, the aim of the study is to systematically manipulate the numbers of effectors (1 or 2 hands combined with 1, 2 or 4 fingers) used to perform the FTT and to investigate the impact of this variation on motor memory consolidation. In a series of seven experiments, we investigate five variants of the tasks and replicate the findings of the two most critical experiments. We hypothesise that the use of more effectors leads to greater post-sleep enhancement relative to post-wake enhancement.

## 2 Material and Methods

### 2.1 Participants

Altogether 144 sport science students participated in the experiment (age = 23.4 years, *SD* = 2.4, 67 = female). Sample size was determined based on previous research of the classic FTT (Walker et al., 2002; Walker, Brakefield, Seidman, et al., 2003). In Experiment 1, 2, 3, 6 and 7, the 120 participants (age = 24.4 years, *SD* = 2.3, 57 = female) were from the Institute of Sports and Sports Sciences at the Heidelberg University (Germany). In Experiment 2 and 5, the 24 participants (age = 21.1 years, *SD* = 3.4, 10 = female) were from the Institute of Sport Science at the University of Bern (Switzerland). Experiment 2 and 5 served as replications from Experiment 1 and 4. Furthermore, participants in Experiment 2 and 5 were the same, thus, strictly speaking, the participants in these two experiments represent within-subject data. Written informed consent was obtained from all the participants. The 7 experiments were carried out in accordance with the 1964 Declaration of Helsinki.

### 2.2 Procedure

Participants visited the laboratory three times (Acquisition, Retention 1, and Retention 2). The participants in the EME groups visited the laboratory for *acquisition* at 8.00 PM (**E**vening), for *retention 1* at 8.00 AM (**M**orning) and for *retention 2* at 8.00 PM (**E**vening). Therefore, in EME groups, a nocturnal sleep interval occurred in the first phase while a wake interval occurred in the second phase of the experiment. The participants in the MEM groups visited the laboratory for *acquisition* at 8.00 AM (**M**orning), for *retention 1* at 8.00 PM (**E**vening) and for *retention 2* at 8.00 AM (**M**orning). Therefore, in the MEM groups, a wake interval occurred in the first phase while a sleep interval occurred in the second phase of the experiment. Participants slept in their own home settings during the night and were instructed not to sleep during the day. No restrictions on caffeine or alcohol intake before or during the wake interval were made.

Participants had to perform a five-digit sequence (4-1-3-2-4) that was shown on a screen in front of them; the instructions were to type the sequence as fast and accurate as possible on a standard computer keyboard (Walker et al., 2002; Walker, Brakefield, Seidman, et al., 2003). The learning phase consisted of 12 learning blocks, each with a duration of 30 seconds. After every learning block participants had a 30 second break, resulting in an overall duration of the learning phase of 12 minutes. Retest 1 and Retest 2 each consisted of two blocks with a 30 second break between the two blocks. This procedure was used in all seven experiments. The respective experiments only differed in the number of hands and fingers used to perform the sequence. In Experiment 1 and 2, the sequence was typed with the index finger of the left hand. In Experiment 3, the index and middle finger of the left hand was used. In Experiment 4 and 5, the classic FTT was performed with the index, middle, ring and pinky finger of the left hand. In Experiment 6, the index finger of both hands was used. While in Experiment 7, the index and middle finger of both hands was used.

### 2.3 Data analysis

The dependent variable in all experiments was the number of correct sequences (CS) per block. To quantify the progress over the course of the learning phase, the number of in the first block of the learning phase (block 1) was compared with that in the last block (block 12) of the learning phase. These two measures represented the pre-test and post-test performance, respectively. The mean number of correct sequences in the last two blocks of the learning phase (block 11 and 12) were defined as the performance post-learning. Furthermore, the mean of the two blocks of retest 1 and 2 were used as the performance measure at retest 1 or 2, respectively. The performance change over sleep (post-sleep gain, PSG) was calculated by subtracting the before sleep from the number of after sleep for each group separately (EME: at retest 1 minus at post-test; MEM: at retest 2 minus at retest 1). The performance change over wake (post-wake gain, pwg) was calculated by subtracting the number of before wake from the number of after wake (EME: at retest 2 minus at retest 1; MEM: at retest 1 minus at post-test). The mean difference between the PSG and the PWG, with consideration of the standard deviation, can be interpreted as the relative sleep gain (RSG), as used in previous publications (Pan & Rickard, 2015; Schmid et al., 2020). The direction of the effect is such that a higher number of and a positive effect represent better performance.

### 2.4 Statistics

To compare the post-sleep gain and the post-wake gain of each group, two-tailed and paired t-tests were used. Cohen’s *d* was used as the effect size and interpreted according to Cohen (1988), with small (≥ 0.2), medium (≥ 0.5) and large (≥ 0.8) effects. A significance level of *p* < .05 was used for all the inferential statistics. All statistical analyses were performed in R (v 3.6.1 with RStudio (v 1.3.959).

## 3 Results

### 3.1 Practice-dependent learning

The number of in all the experiments from block 1 to block 12 is shown in Table 1. Every group in all seven experiments had highly significant performance changes (all p’s < .001) with large effect sizes ranging from *d =* 1.08 to *d =* 2.78 over the course of the learning phase.

**Table 1.** Number of for all the experiments from block 1 to block 12 with the corresponding inferential statistics.

| Experiment | Number of correct sequences at learning phase (SD) |  | Inferential statistics |  |  |
| --- | --- | --- | --- | --- | --- |
|  | Block 1 | Block 12 | Practice dependent learning |  |  |
|  |  |  | *t*(11) | *p* | *d* |
| Exp. 1: 1 Hand with 1 Finger |  |  |  |  |  |
| EME | 13.58 (6.07) | 24.00 (2.98) | 6.46 | < .001 | 1.87 |
| MEM | 13.63 (3.95) | 23.08 (3.26) | 8.54 | < .001 | 2.47 |
| Exp. 2: 1 Hand with 1 Finger replication |  |  |  |  |  |
| EME<sup>3</sup> | 15.22 (3.66) | 24.63 (4.86) | 9.64 | < .001 | 2.78 |
| MEM<sup>3</sup> | 13.08 (4.42) | 21.35 (2.43) | 5.68 | < .001 | 1.64 |
| Exp. 3: 1 Hand with 2 Fingers |  |  |  |  |  |
| EME | 15.25 (5.28) | 22.92 (5.28) | 7.89 | < .001 | 2.28 |
| MEM | 20.66 (8.56) | 25.92 (6.22) | 3.72 | < .001 | 1.08 |
| Exp. 4: 1 Hand with 4 Fingers |  |  |  |  |  |
| EME | 9.67 (5.02) | 17.67 (4.36) | 6.53 | < .001 | 1.89 |
| MEM | 12.42 (3.68) | 19.72 (4.21) | 8.20 | < .001 | 2.37 |
| Exp. 5: 1 Hand with 4 Fingers replication |  |  |  |  |  |
| EME | 15.80 (6.22) | 24.24 (4.63) | 4.50<sup>2</sup> | < .001 | 1.36 |
| MEM | 19.27 (5.38) | 23.17 (5.10) | 4.52 | < .001 | 1.30 |
| Exp. 6: 2 Hands with 2 Fingers |  |  |  |  |  |
| EME | 19.60 (8.76) | 31.42 (6.33) | 5.34<sup>1</sup> | < .001 | 1.54 |
| MEM | 20.75 (6.03) | 34.42 (5.18) | 6.56 | < .001 | 1.89 |
| Exp. 7: 2 Hands with 4 Fingers |  |  |  |  |  |
| EME | 11.83 (9.45) | 28.08 (7.67) | 6.47 | < .001 | 1.87 |
| MEM | 14.78 (9.69) | 26.00 (7.56) | 9.09<sup>2</sup> | < .001 | 2.74 |

Note, <sup>1</sup>df = 11, <sup>2</sup>df = 10, <sup>3</sup>significant difference at block 12 between the two groups (*t*(22) = 2.09, *p* = 0.048, *d* = 0.85)

### 3.2 Offline learning

The descriptive data from the seven experiments at post-learning, retest 1 and retest 2 with the corresponding inferential statisti is shown in Table 2. An overview over the post-sleep gains and the post-wake gains with the corresponding effect sizes is shown in Figure 1A for the EME and in Figure 1B for the MEM group.

**Table 2.**
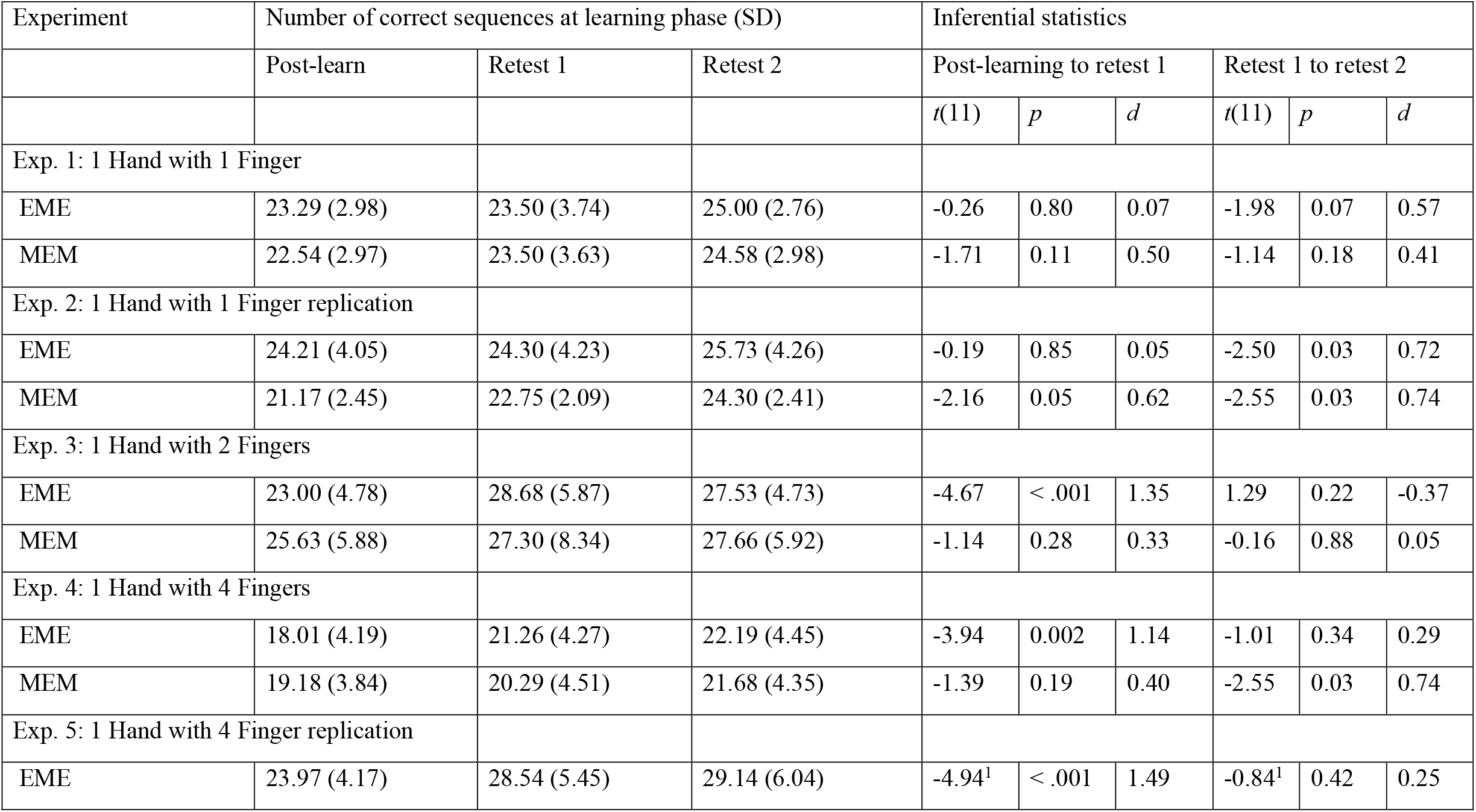
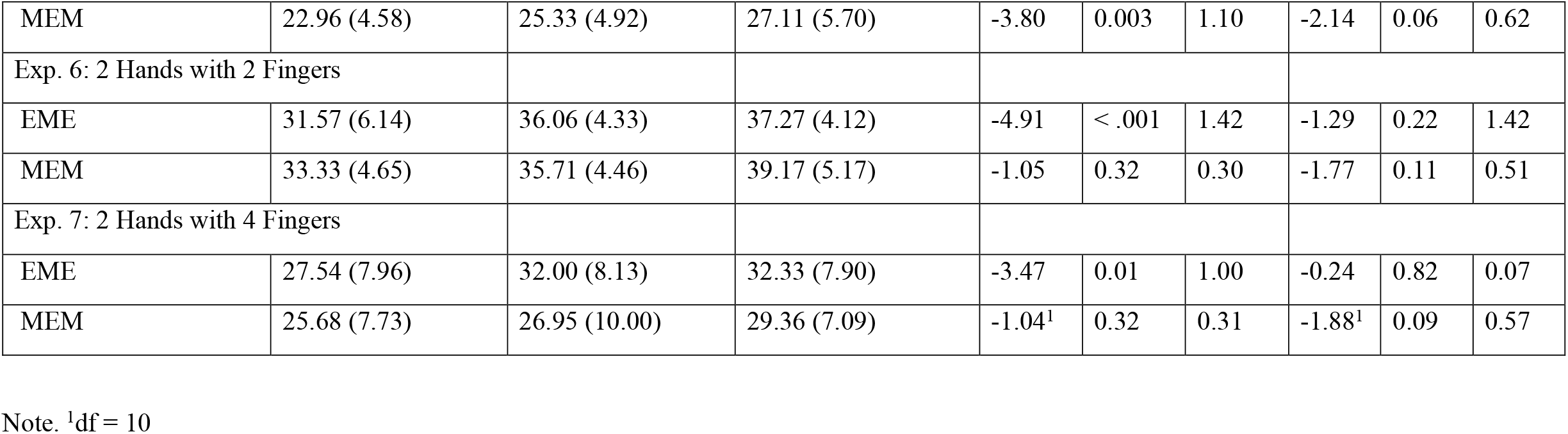
Overview of the number of in all experiments at post-learning, retest 1 and retest 2 with the corresponding inferential statistics.

**Figure 1.**
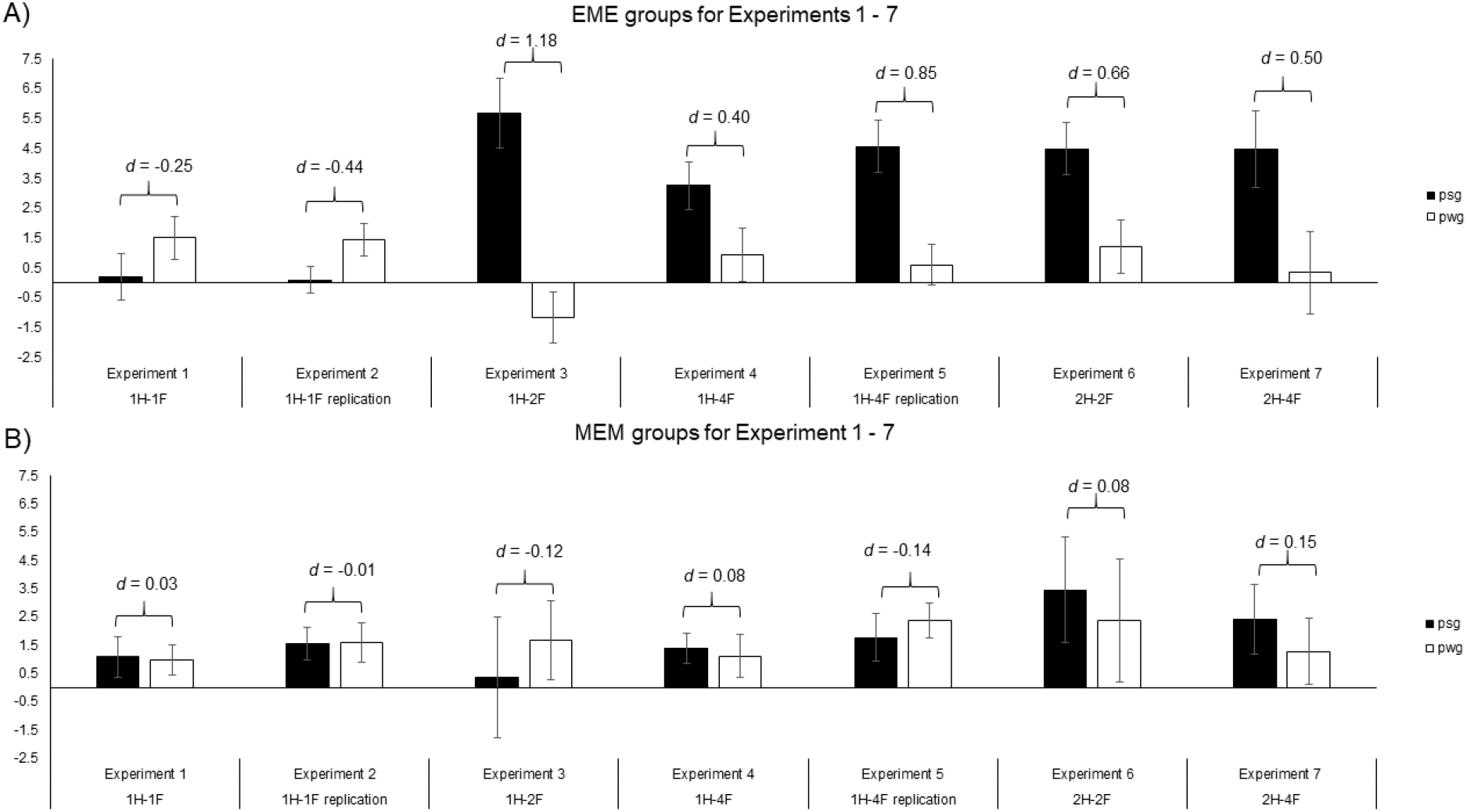
Comparison of the post-sleep gains and the post-wake gains with corresponding effect sizes for all the seven experiments. (A) EME groups (B) MEM groups.

Experiment 1 (1 Hand with 1 Finger). The difference in the post-sleep gain (*M* = 0.21, *SD* = 2.68) and the post-wake gain (*M* = 1.50, *SD* = 2.52) was not significant, *t*(11) = -0.86, *p* = 0.41, *d* = -0.25, in the EME group. No significant differences between the post-sleep gain (*M* = 1.08, *SD* = 2.54) and the post-wake gain (*M* = 0.96, *SD* = 1.85) were found, *t*(11) = 0.10, *p* = 0.92, *d* = 0.03, in the MEM group.

Experiment 2 (1 Hand with 1 Finger replication). The difference in the post-sleep gain (*M* = 0.09, *SD* = 1.60) and the post-wake gain (*M* = 1.43, *SD* = 1.89) was not significant, *t*(11) = - 1.52, *p* = 0.16, *d* = -0.44) in the EME group. No significant differences between the post-sleep gain (*M* = 1.55, *SD* = 2.02) and the post-wake gain (*M* = 1.58, *SD* = 2.43) were found, *t*(11) = -0.04, *p* = 0.97, *d* = -0.01, in the MEM group.

Experiment 3 (1 Hand with 2 Fingers). The difference in the post-sleep gain (*M* = 5.58, *SD* = 4.03) and the post-wake gain (*M* = -1.16, *SD* = 2.98) was significant with a large effect, *t*(11) = 4.09, *p* = 0.002, *d* = 1.18, in the EME group. No significant differences between the post-sleep gain (*M* = 0.36, *SD* = 7.42) and the post-wake gain (*M* = 1.68, *SD* = 4.89) were found, *t*(11) = -0.41, *p* = 0.69, *d* = -0.12, in the MEM group.

Experiment 4 (1 Hand with 4 Fingers). The difference in the post-sleep gain (*M* = 3.25, *SD* = 2.74) and the post-wake gain (*M* = 0.93, *SD* = 3.08) was not significant with a small effect, *t*(11) = 1.38, *p* = 0.20, *d* = 0.40, in the EME group. No significant differences between the post-sleep gain (*M* = 1.38, *SD* = 1.80) and the post-wake gain (*M* = 1.12, *SD* = 2.66) were found, *t*(11) = 0.26, *p* = 0.80, *d* = 0.08, in the MEM group.

Experiment 5 (1 Hand with 4 Finger replication). The difference in the post-sleep gain (*M* = 4.56, *SD* = 2.92) and the post-wake gain (*M* = 0.60, *SD* = 2.26) was significant with a large effect, *t*(10) = 2.82, *p* = 0.02, *d* = 0.85, in the EME group. No significant differences between the post-sleep gain (*M* = 1.78, *SD* = 2.75) and the post-wake gain (*M* = 2.38, *SD* = 2.08) were found, *t*(11) = -0.47, *p* = 0.65, *d* = -0.14, in the MEM group.

Experiment 6 (2 Hands with 2 Fingers). The difference in the post-sleep gain (*M* = 4.49, *SD* = 3.03) and the post-wake gain (*M* = 1.21, *SD* = 3.11) was significant with a medium effect, *t*(11) = 2.27, *p* = 0.04, *d* = 0.66, in the EME group. No significant differences between the post-sleep gain (*M* = 3.46, *SD* = 6.50) and the post-wake gain (*M* = 2.38, *SD* = 7.51) were found, *t*(11) = 0.29, *p* = 0.77, *d* = 0.08, in the MEM group.

Experiment 7 (2 Hands with 4 Fingers). The difference in the post-sleep gain (*M* = 4.46, *SD* = 4.26) and the post-wake gain (*M* = 0.33, *SD* = 4.62) was not significant with a medium sized effect, *t*(11) = 1.72, *p* = 0.11, *d* = 0.50, in the EME group. No significant differences between the post-sleep gain (*M* = 2.41, *SD* = 4.06) and the post-wake gain (*M* = 1.27, *SD* = 3.86) were found, *t*(10) = 0.49, *p* = 0.64, *d* = 0.15, in the MEM group.

## 4 Discussion

The present study aimed to investigate how the number of used effectors to perform a FTT influences performance enhancement after a sleep interval in comparison with a corresponding wake interval. Seven experiments were presented in which the number of effectors used to perform the classic FTT was systematically varied. The five variations included: 1 Hand with 1 Finger, 1 Hand with 2 Fingers, 1 Hand with 4 Fingers, 2 Hands with 2 Fingers, 2 Hands with 4 Fingers. Furthermore, two of the critical experiments were replicated with another set of participants. All five variants of the tasks were learnable, as in all the experiments a large improvement in the number of was observed over the course of the learning phase. Regarding consolidation in the EME group, we report a beneficial effect of sleep in comparison with a corresponding wake interval for all experiments except experimentthose in which the FTT was performed with 1 hand and 1 finger. For the experiments which represents the classic FTT (1 hand with 4 fingers), we replicated previous finding regarding the magnitude of the RSG (Pan & Rickard, 2015; Schmid et al., 2020). However, no specific relationship between the number of effectors used to perform the task and the RSG can be concluded/was identified from Experiments 3, 4/5, 6 und 7, as the effect sizes of these experiments were of the same magnitudexperiment. For the MEM group, we do not report positive findings regarding the effect of sleep in comparison with a wake interval, as no differences between the post-sleep and the post-wake gain have been found.

The current study partially replicates and extends previous findings. Three points should be discussed to integrate the results into the current state of the art: (1) the reported absence of a clear-cut pattern between the numbers of effectors used and the RSG, (2) the time interval between learning and sleep seems to play a role in motor memory consolidation (3) and the potential contributions of electrophysiological features to motor memory consolidation.

(1) The classic FTT has been repeatedly used to look into mechanism of motor memory consolidation and reliably, a beneficial effect of sleep in comparison with a wake interval has been found (Debas et al., 2010; Doyon et al., 2009; Gregory et al., 2014; Nishida & Walker, 2007; Robertson, Pascual-Leone, & Press, 2004; Walker et al., 2002; Walker, Brakefield, Seidman, et al., 2003; Wilson, Baran, Pace-Schott, Ivry, & Spencer, 2012). However, there are few studies that have used variants of the task, like changing the number of effectors used to perform the FTT. Furthermore, two studies that used a relatively gross motor variant of the task, in which the numbers had to be pressed with the whole hand (hand-tapping), reported negative findings. Therefore, in young adults, it seems that sleep does not play a role in the consolidation of hand-tapping in comparison with a corresponding wake interval (Erlacher et al., 2014; Gudberg, Wulff, & Johansen-Berg, 2015). In the study at hand, we could not find a systematic change in the RSG when using a different set of effectors. The performance of the EME group in Experiment 1 shows that there is no relative sleep gain when performing the FTT with only one finger. Sleep even seems to be detrimental to performance; as the pwg is, on a descriptive level, higher that the psg. This finding was replicated in Experiment 2 with a different set of participants. In all the other experiments, the EME groups had relative sleep gains with small to large effect sizes. However, no clear-cut pattern was found to reveal a relationship between the numbers of effectors used to perform the FTT and the RSG.

(2) Recently, the effects of sleep in comparison with a corresponding wake interval have been quantified in two meta-analyses. The results of the EME group in Experiment 4, which represents the classic FTT (Walker et al., 2002; Walker, Brakefield, Seidman, et al., 2003), is in accordance with current meta-analytical evidence regarding this motor task (Pan & Rickard, 2015; Schmid et al., 2020). However, the replication partially failed in the EME group of Experiment 5, such that the effect size is of noticeable difference to that of Experiment 4. This can potentially be explained by the fact that the participants in the replication studies (Experiment 2 and 5) were the same. When beginning experiment 5, participants had already participated in Experiment 2 and thus had some experience with the FTT. Furthermore, we failed to replicate the time-independency that has been shown in earlier finger tapping studies (Walker et al., 2002; Walker, Brakefield, Seidman, et al., 2003), as none of the MEM groups showed significant differences between the post-sleep and post-wake gain. The performed experiments were not designed to test the hypothesis of whether a wake interval between learning and retesting influences the RSG. However, the presented results rather suggest that the placement of sleep immediately after the learning phase is beneficial and that a longer wake interval between learning and testing does lead to diminished gains (Doyon et al., 2009; Holz et al., 2012). This could be explained with interference during the wake interval (Borragán, Urbain, Schmitz, Mary, & Peigneux, 2015; Friedman & Korman, 2016; Korman et al., 2007; Walker, Brakefield, Hobson, & Stickgold, 2003). Future studies should try to disentangle this finding by systematically manipulating the time interval spent awake between learning and sleep, as this information could also be of interest for sport practice and rehabilitation.

(3) The electrophysiological basis of motor sequence learning has been repeatedly investigated. Findings suggest that stage 2 sleep seems to play an important role in the consolidation of motor sequence learning, as the performance improvement overnight correlates with the amount of stage 2 sleep (Bottary, Sonni, Wright, & Spencer, 2016; Korman et al., 2007; Nishida & Walker, 2007; Walker et al., 2002). Furthermore, the frequency of ? sleep spindles – a key feature of stage 2 sleep – appears to correlate with performance improvement overnight (Barakat et al., 2013; Nishida & Walker, 2007; Tucker & Fishbein, 2009). Although we did not find a systematic change in the relative sleep gain when the FTT is performed with a different number of effectors, polysomnographic record might give further insights into the nature of motor memories and their? consolidation. As no polysomnographic record was performed in this study, future studies should also try to find differences in the consolidation of motor tasks performed with a different number of effectors.

## 5 Conclusion

The presented results suggest that the numbers of effectors used to perform a FTT does not predict the RSG. Furthermore, the time interval between learning and sleep seems to be of importance to the beneficial effect of sleep on motor memory consolidation. Future studies should aim to experimentally manipulate the time window between learning and testing and to look into the electrophysiological basis of performing a FTT with a different set of effectors.

